# UCHL3 regulates subgenomic flaviviral RNA condensates to promote virus propagation

**DOI:** 10.1101/2025.09.05.674517

**Authors:** Oscar Trejo-Cerro, Anna Beekmayer-Dhillon, Qi Wen Teo, Lewis Siu, Ming Yuan Li, Sumana Sanyal

## Abstract

In this study we demonstrate a previously uncharacterised post-translational regulatory mechanism governing flavivirus replication through the deubiquitylating enzyme ubiquitin C-terminal hydrolase L3 (UCHL3). Using activity-based protein profiling, we identified UCHL3 as a key cellular factor activated during Zika virus (ZIKV) and dengue virus (DENV) infections. CRISPR-Cas9 knockout experiments demonstrated that UCHL3 deficiency impairs flavivirus replication and viral protein expression across multiple cellular models. The underlying molecular mechanism involves UCHL3-mediated stabilisation of subgenomic flaviviral RNA (sfRNA)-containing biomolecular condensates. Through biotinylated sfRNA-interactome capture assays, we show that UCHL3 physically interacts with sfRNA-containing ribonucleoprotein complexes alongside G3BP1. Importantly, UCHL3 depletion triggers inappropriate RNase L activation, leading to sfRNA relocalisation from protective P-bodies to degradative compartments, such as RNase L-induced bodies (RLBs) as reported previously, resulting in viral RNA decay. Our rescue experiments confirmed that RNase L knockdown restores viral replication in UCHL3-deficient cells. This pro-viral effect of UCHL3 operates through interferon-independent mechanisms, as demonstrated by persistent replication defects even upon exogenous interferon treatment. This work therefore identifies UCHL3 as a molecular switch controlling the balance between pro-viral and antiviral RNA condensates, representing a promising host dependency factor for broad-spectrum flavivirus intervention strategies.

---

Flaviviruses, such as Zika virus (ZIKV), dengue virus (DENV), and West Nile virus (WNV), pose significant global health threats on account of their ability to cause severe neurological complications, haemorrhagic fever, and birth defects^1^. These positive-sense RNA viruses have evolved intricate strategies to hijack cellular machinery for efficient replication while evading innate immune responses^2^. A key feature of flavivirus transmissibility and their epidemic potential is the generation of subgenomic flavivirus RNA (sfRNA), a highly structured noncoding RNA produced as a result of incomplete viral genomic RNA degradation by the host 5’-3’ exoribonuclease XRN1^3,4^. This conserved feature across the flavivirus family serves as a virulence determinant. Multiple lines of evidence show that viruses unable to produce sfRNAs replicate less efficiently, are less pathogenic in animal models, and cause reduced cytotoxicity in cell culture^5,6^.

Emerging evidence has indicated several potential functions of sfRNA in both the human and vector hosts. These include antagonism of antiviral immunity, promotion of replication, viral fitness and host adaptation^5,6^. These molecules are thought to function as molecular decoys that sequester RNA-binding proteins and interfere with antiviral signalling pathways^7^. For example, sfRNAs have been shown to mimic natural substrates for Dicer and Argonaute, interfering with antiviral RNAi in insect cells^8^. Recent evidence has revealed that sfRNAs organise into dynamic ribonucleoprotein complexes resembling stress granules, forming biomolecular condensates that concentrate viral and cellular factors essential for replication and immune evasion^9^. The sfRNA-containing condensates appear to be distinct from classical stress granules: they are smaller in size (∼1µm), and colocalise with G3BP1, PABPC1, UBAP2L, and poly(A) RNA^9^.

The formation, composition, and dynamics of biomolecular condensates have been reported to be tightly regulated by post-translational modifications, particularly ubiquitylation^10,11^. The ubiquitin-proteasome system controls protein stability, localisation, signalling, and protein-protein interactions within these condensed phases^12,13^. While ubiquitin conjugation can target proteins for various processes such as proteasomal degradation, immune signalling or autophagy, deubiquitylating enzymes (DUBs) reverse these modifications to stabilise proteins and modulate signalling cascades^14^. Many viruses exploit enzymes of ubiquitylation and ubiquitin like modifications and their corresponding hydrolases to enhance replication, transmission and immune evasion^15,16^, with several known examples of viral proteins encoding DUB activities or recruiting cellular DUBs to modify host factors in favour of virus dissemination^17,18^.

Ubiquitin C-terminal hydrolase L3 (UCHL3) is a cysteine protease that belongs to the ubiquitin C-terminal hydrolase (UCH) family of deubiquitylating enzymes^19^. UCHL3 cleaves ubiquitin from small adducts and processes polyubiquitin chains, thereby regulating protein stability and cellular signalling. While UCHL3 has been implicated in various cellular processes including DNA repair^20^, cell cycle progression^21^, and stress responses^22^, its role in viral infection remains poorly understood. The enzyme’s ability to modulate protein stability through deubiquitylation makes it an attractive target for viral manipulation, particularly in the context of RNA granule dynamics where protein turnover critically influences condensate composition and function.

Despite growing appreciation for the importance of sfRNA condensates in flavivirus biology, the molecular mechanisms governing their assembly, maintenance, and regulation remain poorly understood. The post-translational control of condensate-associated proteins represents a largely unexplored regulatory layer that could significantly impact on immune evasion, viral persistence and spread. Given the central role of ubiquitylation in controlling protein fate and the documented ability of viruses to hijack deubiquitylating enzymes, we hypothesised that cellular DUBs might regulate sfRNA condensate dynamics to promote flavivirus replication.

Here, we perform activity-based protein profiling to identify UCHL3 as specifically activated during infection. We investigate the role of UCHL3 in flavivirus infection and demonstrate that this deubiquitylating enzyme functions as a key pro-viral factor. We show that UCHL3 is induced upon infection, partially translocates to the nucleus, and potentially associates with biomolecular condensates retaining sfRNA within them. Loss of UCHL3 results in relocalisation of sfRNA away from P-bodies to possibly RNAse L-induced bodies (RLBs), resulting in degradation of viral genomic RNA. Our findings therefore reveal that UCHL3 prevents sfRNA sequestration into antiviral RLBs by stabilising XRN1 condensate scaffold proteins. This strategy maintains sfRNA in protective P-body compartments, thereby promoting viral genome stability and efficient spread. These results establish a novel post-translational regulatory mechanism controlling the balance between pro-viral and antiviral RNA condensates, and identify UCHL3 as a promising therapeutic target for broad-spectrum flavivirus intervention.

## Results

### UCHL3 is activated during viral infection and required for flavivirus production and spread

To identify deubiquitylases (DUBs) that are activated during viral infection, we employed activity-based protein profiling using the ubiquitin-vinyl methyl ester (Ub-VME) probe as reported previously^17^, which covalently labels catalytically active DUBs (**Figure 1A**). Following infection with influenza^17^, Zika and dengue viruses, we observed specific activation of several DUBs, including UCHL3, a cytoplasmic deubiquitylase of the ubiquitin C-terminal hydrolase family (**Figure 1A**). This activation pattern suggested a conserved role for this enzyme across different viral families. To validate and quantify this activation, we first measured UCHL3 protein levels over infection time course. Immunoblot analyses demonstrated that UCHL3 undergoes upregulation in response to Zika infection, with protein levels increasing substantially above basal expression in both Huh7 and A549 cells (**Figure 1B**).

**Figure 1.**
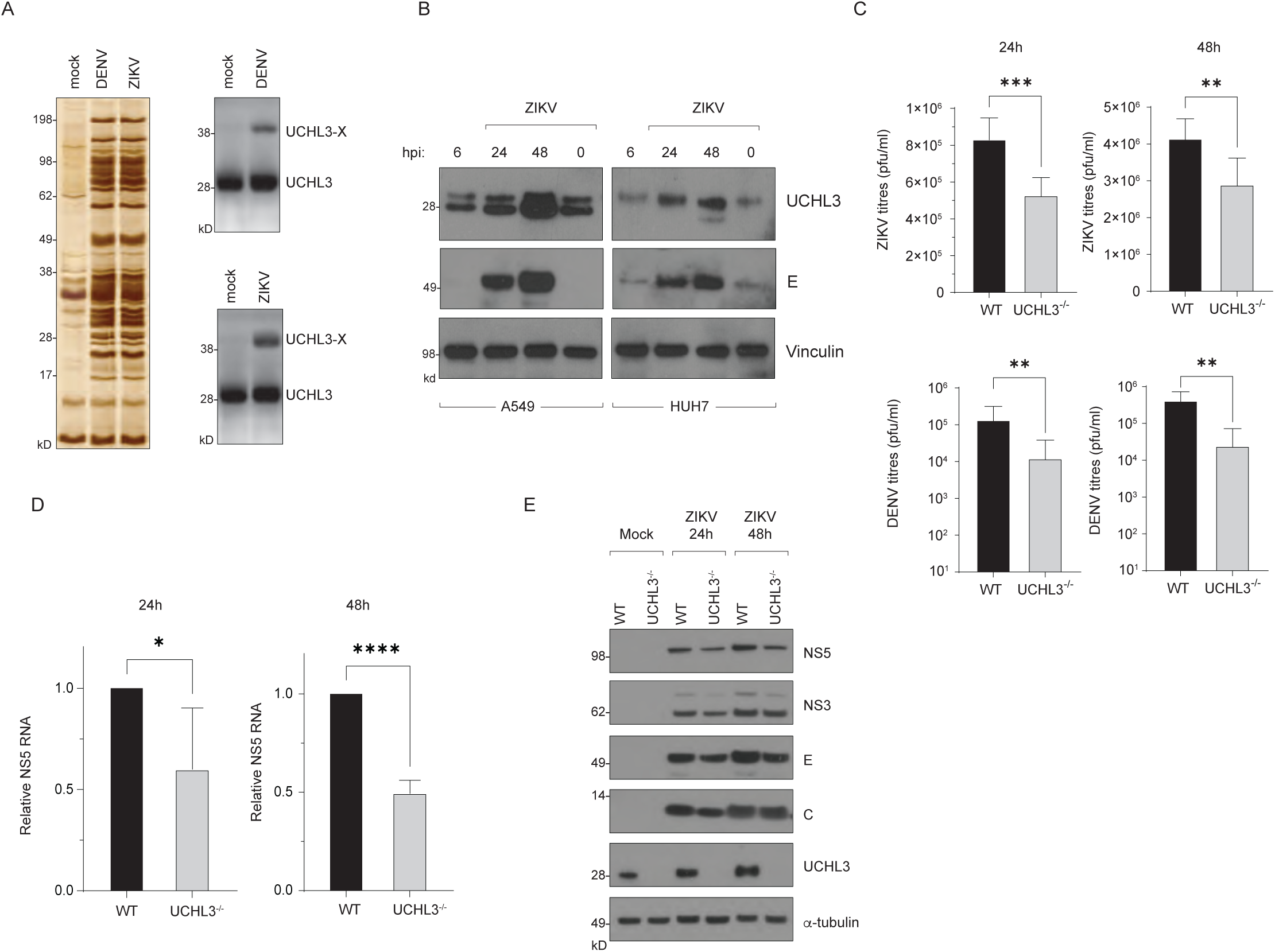
UCHL3 is activated during viral infection and required for flavivirus production and spread. **(A)** Activity-based protein profiling using ubiquitin-vinyl methyl ester (Ub-VME) to capture deubiquitylating enzymes activated during viral infection. *Left panel*: silver stain of total protein profiles captured; *right panel*: immunoblot detection of UCHL3 and (UCHL3-Ub) in mock-infected, dengue virus (DENV)-, and Zika virus (ZIKV)-infected cells. **(B)** Time-course analysis of UCHL3 protein expression during ZIKV infection. Immunoblot analysis in A549 and Huh7 cell lines of UCHL3 upregulation at 6, 24, and 48 hours post-infection (hpi) compared to uninfected controls (0 hpi). Viral envelope (E) protein and vinculin (loading control) are shown. **(C)** UCHL3 deficiency impairs flavivirus replication. Viral titres (pfu/ml) in wild-type (WT) and UCHL3^-/-^ A549 cells infected with ZIKV or DENV at 24 and 48 hours post-infection. Data represent mean ± SD from triplicate experiments. **P < 0.01, ***P < 0.001. **(D)** Reduced viral RNA accumulation in UCHL3^-/-^ cells. Quantitative RT-PCR analysis of NS5 RNA levels normalised to actin in WT and UCHL3^-/-^ cells at 24 and 48 hours post-infection. Data represent mean ± SD. *P < 0.05, ****P < 0.0001. **(E)** Diminished viral protein expression in UCHL3^-/-^ cells. Immunoblot analysis of viral non-structural proteins (NS5, NS3), structural proteins (E, C), and UCHL3 in WT and UCHL3^-/-^ A549 cells under mock infection or ZIKV infection conditions at 24 and 48 hours post-infection. α-tubulin serves as loading control.

To investigate the functional significance of UCHL3 in flavivirus infection, we generated UCHL3 knockout (KO) A549 cells using CRISPR-Cas9 gene editing (**supplementary Figure S1A**) and infected them with either Zika or dengue virus. UCHL3 deletion significantly reduced replication of both flaviviruses, with a more significant reduction of viral titres in dengue, both 24 and 48 hours post-infection compared to wild-type cells (**Figure 1C**). This reduction in viral output was accompanied by a corresponding decrease in viral RNA accumulation, as measured by NS5 RNA levels, which showed progressive loss over time in UCHL3 KO cells infected with either virus (**Figure 1D**). In contrast, the impact of UCHL3 deficiency on influenza virus replication was opposite of the flavivirus phenotype where UCHL3^-/-^ cells displayed an increase in influenza polymerase activity, with no significant impact on infectious virus production (**supplementary figure S1B, S1C**).

The impaired replication in UCHL3-deficient cells was further confirmed by analysis of viral protein expression. Western blot analysis revealed a modest reduction in steady state levels of multiple viral proteins, including the non-structural proteins NS5 and NS3, as well as the structural protein E, in UCHL3 KO cells compared to wild-type controls (**Figure 1E**). This broad reduction in viral protein accumulation across both flaviviruses suggests that UCHL3 affects either viral RNA translation or replication at a fundamental level rather than targeting specific viral proteins, consistent with its conserved pro-viral role.

### UCHL3 undergoes subcellular redistribution upon infection, facilitating flavivirus propagation

To characterise UCHL3 in virus infected cells, we performed subcellular localisation analyses in mock and infected cells. Specificity of UCHL3 staining was validated in wild-type and UCHL3^-/-^ cells (**supplementary Figure S2**). Immunofluorescence analyses revealed striking changes in UCHL3 distribution following viral infection. In mock-infected cells, UCHL3 displayed predominantly cytoplasmic localisation. However, ZIKV infection triggered substantial nuclear accumulation of UCHL3 whilst maintaining its cytoplasmic presence (**Figure 2A**). Quantitative analysis confirmed that this dual-compartment localisation occurred in both productively infected cells and the bystander population (**Figure 2B**). Biochemical fractionation studies corroborated the microscopy findings, indicating clear redistribution of UCHL3 between cytoplasmic and nuclear compartments during infection (**Figure 2C**). The fact that both infected and bystander cells displayed this response suggests that UCHL3 relocalisation is triggered by soluble inflammatory mediators or cellular stress responses rather than requiring direct contact with viral proteins. Importantly, UCHL3 did not colocalise with viral double-stranded RNA (dsRNA) replication intermediates, suggesting that it does not directly associate with viral replication complexes.

**Figure 2.**
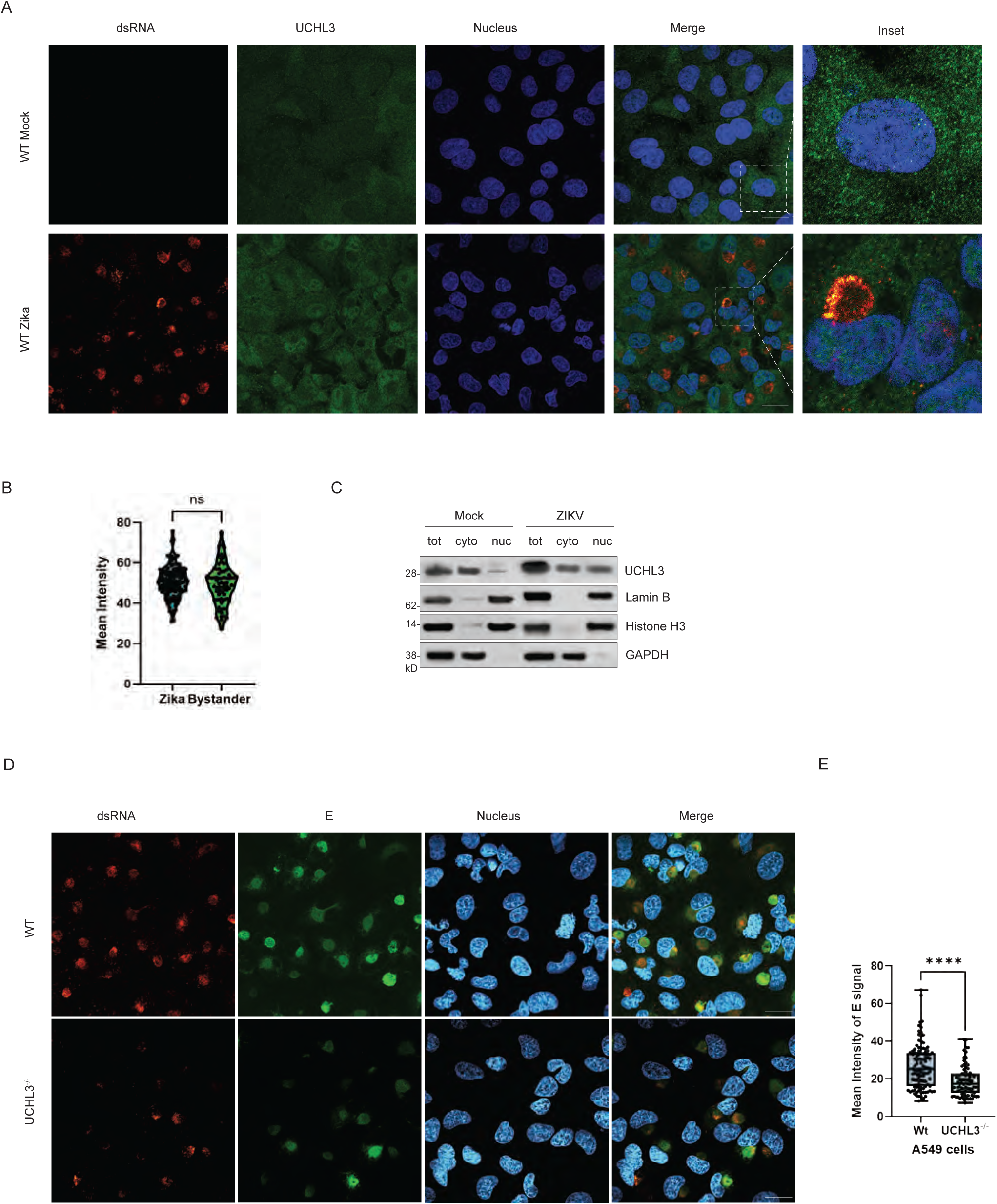
UCHL3 undergoes subcellular redistribution upon infection, facilitating flavivirus propagation. **(A)** Confocal microscopy images of WT A549 cells under mock infection (upper panel) and Zika virus infection at 24 hours post-infection (lower panel) and stained for double-stranded viral RNA (dsRNA, red) to mark replication intermediates, UCHL3 (green), and nuclei (DAPI, blue). Merged images demonstrate UCHL3 redistribution from predominantly cytoplasmic localisation in mock-infected cells to dual cytoplasmic-nuclear distribution during infection. Inset panels provide higher magnification views of individual cells. Scale bar represents 20 μm. **(B)** Quantitative analysis of UCHL3 nuclear accumulation in infected versus bystander cell populations. Violin plot shows mean fluorescence intensity measurements for both productively infected (Zika) and neighbouring uninfected (Bystander) cells. ns, not significant. **(C)** Biochemical validation of UCHL3 subcellular redistribution. Cell fractionation followed by immunoblot analysis of total cellular lysates (tot), cytoplasmic (cyto), and nuclear (nuc) fractions from mock-infected and ZIKV-infected A549 cells. Lamin B (nuclear marker), Histone H3 (nuclear marker), and GAPDH (cytoplasmic marker) serve as fractionation controls. **(D)** Immunofluorescence analysis of WT and UCHL3^-/-^ A549 cells infected with ZIKV. dsRNA (red) and viral envelope protein E (green) mark replication and assembly sites, respectively. UCHL3-deficient cells exhibit diminished and disorganised viral compartments compared to structured perinuclear replication organelles in WT cells. Scale bar represents 20 μm. **(E)** Violin plot analysis of E protein mean fluorescence intensity in WT versus UCHL3^-/-^ A549 cells showing significant reduction in viral protein levels in knockout cells. Data represent measurements from multiple fields across 3 independent experiments. ****P < 0.0001.

To assess the functional consequences of UCHL3 deficiency on virus-induced ER remodelling, we examined the organisation of viral replication and assembly sites using dsRNA and E protein as markers. In wild-type cells, these structures typically manifest as distinct perinuclear compartments representing organised replication organelles and assembly sites. Consistent with the observed reduction in viral titres and RNA levels, UCHL3-deficient cells exhibited significant disruption of these virus-induced compartments (**Figure 2D**), with marked reduction in the intensity of viral protein and dsRNA accumulation (**Figure 2E**). This architectural defect suggests that UCHL3 contributes to establishing or maintenance of productive viral replication/assembly environments.

These findings collectively demonstrate that UCHL3 functions as a pro-viral host factor that undergoes selective activation and relocalisation during flavivirus infection. The infection-induced nuclear accumulation of this normally cytoplasmic deubiquitylase, combined with its conserved requirement across multiple flaviviruses, suggests that UCHL3 serves distinct but complementary functions in different cellular compartments during the viral life cycle.

### UCHL3 associates with flaviviral sfRNA

To assess the role of UCHL3 in Zika virus infection, we overexpressed ectopic Flag-HA-UCHL3 (**Supplementary Figure S3A**). Immunoblotting analyses confirmed expression of the Flag-HA-UCHL3 construct (**Supplementary Figure S3A**). Overexpression of UCHL3 resulted in increased levels of viral E and NS5 proteins. Quantitative assessment of viral replication revealed that ectopic UCHL3 expression moderately enhanced Zika virus propagation in a time-dependent manner (**Supplementary Figure S3B**). At 24 hours post-infection, UCHL3 overexpression resulted in a modest but measurable increase in viral titres. By 48 hours post-infection this effect was more pronounced.

To elucidate the molecular basis of UCHL3’s pro-viral function, we investigated whether UCHL3 directly interacts with viral components during flavivirus infection. Co-immunoprecipitation experiments in cells transfected with Flag-HA-UCHL3 and infected with Zika virus did not reveal any association between UCHL3 and viral structural or non-structural proteins (**Figure 3A**).

**Figure 3.**
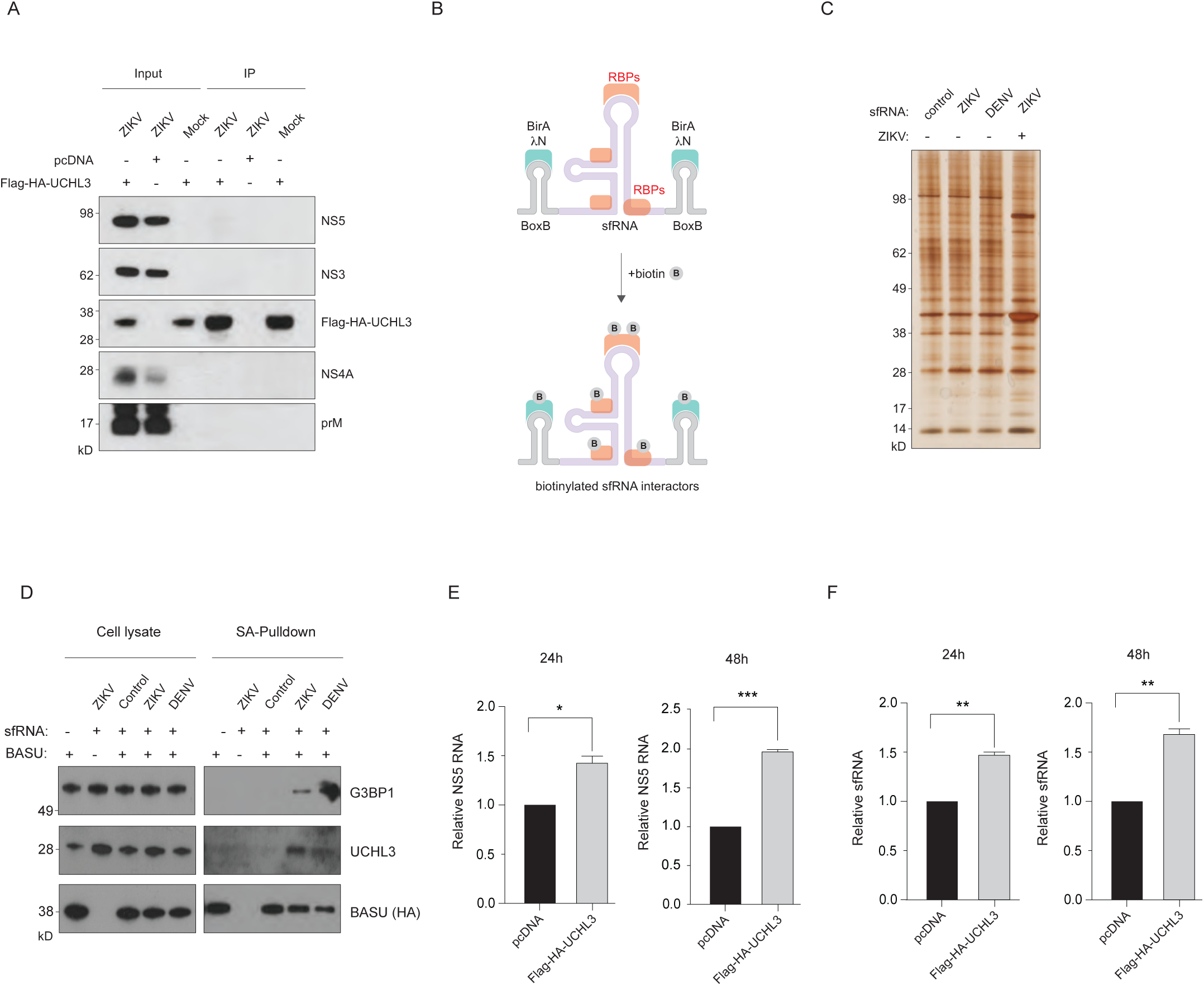
UCHL3 associates with flaviviral sfRNA. **(A)** Co-immunoprecipitation analysis of Flag-HA-UCHL3 from 293 cells transfected with either empty vector control (pcDNA3.1) or Flag-HA-UCHL3 construct and infected with ZIKV or mock-infected. Input lysates and immunoprecipitated (IP) fractions were analysed by immunoblotting for viral non-structural proteins NS5, NS3, and NS4A, viral structural protein prM, and Flag-HA-UCHL3. **(B)** Schematic representation of the biotinylated sfRNA-interactome capture strategy. The engineered sfRNA construct contains flanking BoxB stem-loop structures that specifically recruit the BirA-λN fusion protein (BASU). Upon biotin supplementation, BASU biotinylates proteins in close proximity to sfRNA, enabling streptavidin-mediated affinity purification of sfRNA-associated ribonucleoprotein complexes. **(C)** Silver staining analysis of sfRNA-interacting protein complexes. Total protein profiles from streptavidin pulldowns using control RNA motif, ZIKV sfRNA, or DENV sfRNA constructs in the presence or absence of ZIKV infection for infection-dependent recruitment of cellular factors to sfRNA assemblies. **(D)** Streptavidin-affinity capture followed by immunoblot analysis showing selective enrichment of UCHL3 and G3BP1 in the presence of ZIKV or DENV sfRNA constructs. Cell lysates and streptavidin pulldown (SA-Pulldown) fractions show specific association of these proteins with sfRNA-containing complexes. BASU (HA-tagged) serves as an internal control for the capture system. **(E-F)** Quantitative RT-PCR analysis of (E) NS5 genomic RNA and (F) sfRNA levels in cells transfected with either pcDNA3.1 empty vector or Flag-HA-UCHL3 at 24 and 48 hours post-ZIKV infection. Ectopic UCHL3 expression increases both viral genomic and subgenomic RNA species compared to vector controls. Data represent mean ± SD from triplicate experiments. *P < 0.05, **P < 0.01, ***P < 0.001.

Given the potential for RNA binding by UCHL3, we developed a biotinylated sfRNA-interactome capture strategy to systematically identify sfRNA-interacting proteins (**Figure 3B**) from cells expressing the engineered sfRNA. Silver staining analysis of biotinylated sfRNA-interactors captured on streptavidin beads revealed distinct protein profiles between control and Zika-infected conditions (**Figure 3C**), indicating infection-dependent recruitment of cellular factors to sfRNA complexes.

Streptavidin-affinity capture coupled with immunoblot analysis demonstrated that UCHL3 is specifically recruited to sfRNA ribonucleoprotein complexes (**Figure 3D**). In the presence of sfRNA from either Zika virus or dengue virus, UCHL3 was efficiently captured in the streptavidin pulldown fraction, while control conditions lacking sfRNA showed minimal UCHL3 recovery. Importantly, this sfRNA-mediated recruitment of UCHL3 was accompanied by the co-purification of G3BP1, a well-characterised stress granule and P-body component known to associate with sfRNA. The parallel enrichment of UCHL3, G3BP1, and the biotinylated internal control (BASU) in the pulldown fraction indicates that UCHL3 is incorporated into sfRNA-containing ribonucleoprotein assemblies.

To assess the functional consequences of UCHL3’s association with viral RNA complexes, we performed quantitative RT-PCR analysis of viral RNA levels in cells expressing UCHL3 (**Figures 3E-F**). Ectopic expression of Flag-HA-UCHL3 resulted in significant increase of NS5 genomic RNA accumulation at both 24 hours and 48 hours post-infection. Similarly, sfRNA levels were substantially elevated in UCHL3-overexpressing cells. These data indicate that UCHL3 not only physically associates with viral RNA-protein complexes but also functionally promotes the accumulation of both genomic and subgenomic viral RNA species. Collectively, these findings suggest that UCHL3 might function to regulate sfRNA stability and accumulation during flavivirus infection.

### UCHL3 deficiency triggers RNase L activation and sfRNA sequestration

To elucidate the molecular mechanism by which UCHL3 depletion impairs flavivirus replication, we investigated its impact on the 2’-5’ oligoadenylate synthetase/RNase L (OAS/RNase L), a key innate immune effector system activated by viral dsRNA. Quantitative RT-PCR analysis of 2’-5’ linked oligoadenylates (2-5A) levels revealed a significant accumulation of these endogenous RNAse L activators in UCHL3^-/-^ cells compared to wild-type controls during Zika virus infection (**Figure 4A**). This enhanced 2-5A synthesis was accompanied by pronounced ribosomal RNA (rRNA) degradation, as evidenced by electrophoretic analysis demonstrating characteristic rRNA cleavage products in UCHL3^-/-^ cells (**Figure 4B**). The appearance of rRNA cleavage bands below the intact 28S and 18S ribosomal species provides compelling biochemical evidence for enhanced ribonuclease activity in the absence of UCHL3.

**Figure 4.**
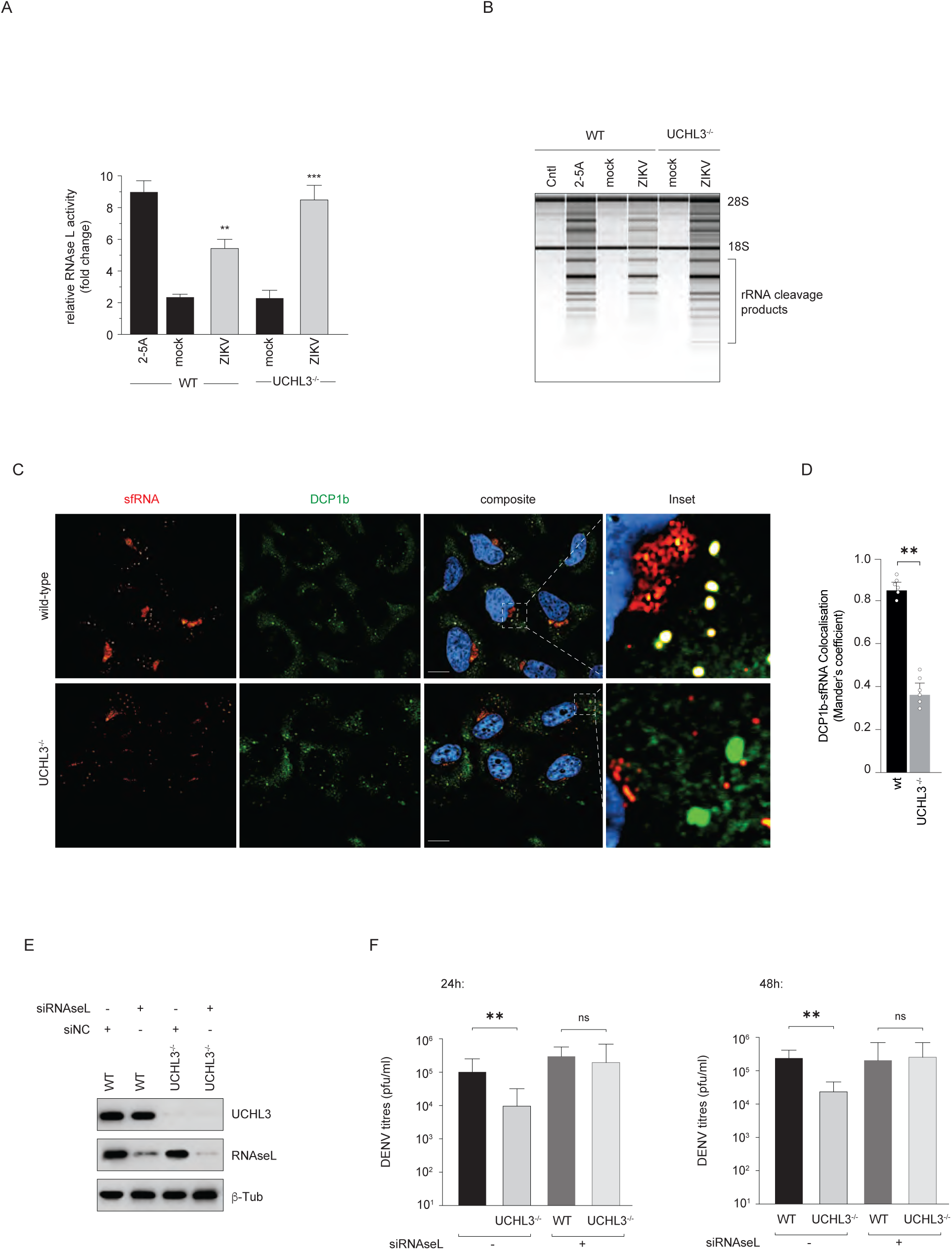
UCHL3 deficiency triggers RNase L activation and sfRNA sequestration. **(A)** Quantitative analysis of RNase L activation in UCHL3^-/-^ cells compared to wild-type (WT) controls during ZIKV infection. 2-5A treatment serves as a positive control for RNase L pathway activation. Data represent mean ± SD from triplicate experiments. **P < 0.01, ***P < 0.001. **(B)** Electrophoretic analysis of total RNA extracted from WT and UCHL3^-/-^ A549 cells under mock infection, ZIKV infection, or 2-5A treatment conditions. UCHL3^-/-^ cells exhibit characteristic rRNA cleavage products below the intact 28S and 18S ribosomal species, providing biochemical evidence for hyperactive ribonuclease activity in the absence of UCHL3. **(C)** sfRNA relocalises from P-bodies to antiviral condensates in UCHL3-deficient cells. Fluorescence in situ hybridisation coupled with immunofluorescence microscopy reveals sfRNA subcellular distribution (red) relative to P-body marker DCP1b (green) in WT and UCHL3^-/-^ cells infected with ZIKV. In WT cells, sfRNA colocalises with DCP1b-positive P-bodies, whilst in UCHL3 UCHL3^-/-^ cells, sfRNA redistributes to distinct cytoplasmic structures consistent with RNase L-induced bodies (RLBs). Composite images show DAPI-stained nuclei (blue). Inset panels provide high-magnification views of sfRNA-containing structures. Scale bar represents 10 μm. **(D)** Quantitative colocalisation analysis demonstrates reduced sfRNA-P-body association in UCHL3-deficient cells. Manders coefficient analysis measuring DCP1b-sfRNA colocalisation shows significant reduction in sfRNA retention within protective P-body compartments in UCHL3^-/-^ cells compared to WT controls. Data represent measurements from multiple fields across 3 independent experiments. **P < 0.01. **(E)** Immunoblot analysis for depletion of RNase L protein levels following siRNA treatment compared to non-targeting control (siNC) in both WT and UCHL3^-/-^ A549 cells. UCHL3 levels remain unaffected by RNase L knockdown. β-tubulin serves as loading control. **(F)** DENV viral titres (pfu/ml) in WT and UCHL3^-/-^ cells treated with control siRNA (siNC) or RNase L-targeting siRNA (siRNaseL) at 24 and 48 hours post-infection. RNase L knockdown rescues viral replication defects in UCHL3^-/-^ cells, demonstrating that impaired viral production results from inappropriate RNase L activation. Data represent mean ± SD from triplicate experiments. **P < 0.01; ns, not significant.

To visualise the spatial consequences of enhanced RNase L activity on viral RNA localisation, we employed fluorescence in situ hybridisation (FISH) coupled with immunofluorescence microscopy to track sfRNA subcellular distribution. In wild-type cells, sfRNA exhibited characteristic colocalisation with P-body markers (DCP1b), consistent with its incorporation into pro-viral ribonucleoprotein condensates (**Figure 4C**). However, in UCHL3^-/-^, sfRNA underwent relocalisation away from DCP1b-positive P-bodies, instead appearing in distinct cytoplasmic structures that likely represent RNase L-induced bodies (RLBs). These antiviral condensates are characterised by enhanced RNase L activity and represent a cellular defence mechanism for viral RNA degradation. Colocalisation analysis demonstrated a significant reduction in sfRNA-P-body association in UCHL3-deficient cells (**Figure 4D**), indicating that UCHL3 is essential for maintaining sfRNA within protective pro-viral compartments.

To establish the causal relationship between RNase L hyperactivation and the observed phenotypes, we performed rescue experiments using RNase L knockdown in the wild-type and UCHL3-deficient background (**Figure 4E**). Western blot analysis confirmed efficient depletion of RNase L protein levels following siRNA treatment in both wild-type and UCHL3^-/-^ cells, whilst UCHL3 levels remained unaffected by the RNase L knockdown. At 24 hours post-infection, UCHL3^-/-^ cells showed reduced dengue virus titres compared to wild-type controls. Remarkably, knockdown of RNase L in UCHL3^-/-^ cells substantially rescued viral replication, with titres similar to wild-type cells. This rescue effect was maintained at 48 hours post-infection, demonstrating that the reduced viral titres in UCHL3 deficient cells is largely RNase L hyperactivation dependent.

These findings collectively demonstrate that UCHL3 functions as a critical regulator that prevents inappropriate RNase L activation during flavivirus infection. In the absence of UCHL3, sfRNA relocalises from protective P-bodies to potentially degradative RLBs, ultimately resulting in enhanced viral RNA decay and reduced viral persistence. This represents an intricate host-pathogen interaction where UCHL3 serves as a molecular switch that tilts the balance between antiviral and pro-viral condensate formation, thereby determining the fate of viral RNA and the success of infection.

### UCHL3 promotes Zika virus infection through interferon-independent mechanisms

To investigate whether UCHL3’s pro-viral function operates through modulation of cellular antiviral responses, we examined interferon-stimulated gene (ISG) expression under various combinations of viral infection and exogenous interferon treatment. This experimental design allows us to dissect the complex interplay between UCHL3, innate immune signalling, and viral replication, to determine whether UCHL3’s effects are primarily immunomodulatory or operate through alternative mechanisms.

First, we analysed ISG15 mRNA levels, a canonical marker of interferon pathway activation, in wild-type and UCHL3^-/-^ cells under four distinct conditions: mock infection, mock infection with exogenous interferon treatment, Zika virus infection alone, and Zika virus infection followed by interferon treatment (**Figure 5A**). In mock-infected conditions, both wild-type and UCHL3^-/-^ cells displayed minimal ISG15 expression as anticipated, whereas exogenous interferon treatment resulted in a dramatic increase in ISG15 in wild-type cells, and a blunted response in UCHL3^-/-^ cells. This indicates that UCHL3 functions as a positive regulator of interferon signalling, likely acting downstream of the interferon receptor to facilitate transcriptional responses. Zika virus infection alone resulted in suppressed ISG15 expression in both cell types, consistent with established flavivirus immune evasion mechanisms. While wild-type cells maintained relatively low ISG15 levels, UCHL3^-/-^ cells exhibited further reduction, indicating that it helps ISG expression even under conditions of viral immune suppression. When Zika virus infection was combined with exogenous interferon treatment, we observed partial restoration of ISG15 expression in wild-type cells, demonstrating that some interferon responsiveness is retained despite viral countermeasures. However, this rescue was significantly impaired in UCHL3^-/-^ cells, indicating that UCHL3 deficiency results in persistent refractoriness to interferon signalling, even when interferon is provided exogenously.

**Figure 5.**
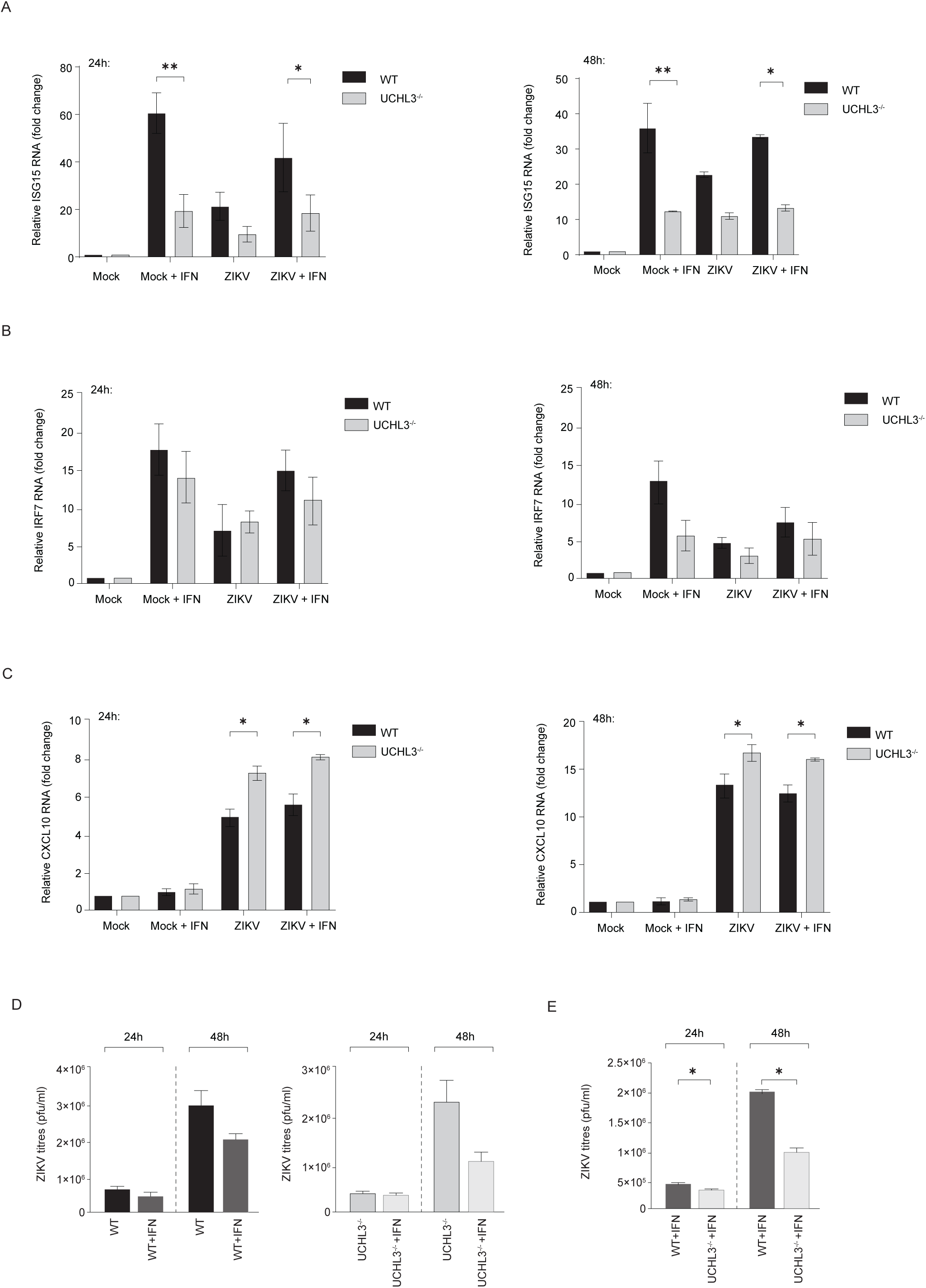
UCHL3 promotes Zika virus infection through interferon-independent mechanisms. **(A)** Quantitative RT-PCR analysis of ISG15 mRNA levels in wild-type (WT) and UCHL3^-/-^ A549 cells under four experimental conditions: mock infection, mock infection with exogenous type-I interferon treatment (Mock + IFN), ZIKV infection alone, and ZIKV infection with interferon treatment (ZIKV + IFN) at 24 and 48 hours. Data represent mean ± SD from triplicate experiments. *P < 0.05, **P < 0.01. **(B)** Differential interferon-stimulated gene responses in UCHL3-deficient cells. IRF7 mRNA expression analysis under identical experimental conditions as (A). **(C)** Paradoxical CXCL10 expression patterns in UCHL3 deficiency. CXCL10 mRNA levels demonstrate contrasting regulation, with UCHL3^-/-^ cells exhibiting enhanced expression during viral infection but reduced responsiveness to interferon treatment, suggesting differential regulatory mechanisms for distinct interferon-responsive genes. *P < 0.05. **(D)** UCHL3’s pro-viral effects persist under interferon-equalised conditions. ZIKV titres (pfu/ml) in WT and UCHL3^-/-^ cells with and without exogenous interferon treatment demonstrate that UCHL3^-/-^ cells maintain significantly reduced viral replication even when interferon responses are bypassed through exogenous supplementation. **(E)** Direct comparison confirms interferon-independent pro-viral activity. ZIKV titres comparing interferon-treated WT and UCHL3^-/-^ cells under identical interferon stimulation conditions demonstrate persistent replication defects in UCHL3-deficient cells. Data represent mean ± SD from triplicate experiments. *P < 0.05.

To determine whether this pattern extends to other ISGs, we examined IRF7 expression under identical conditions (**Figure 5B**). The results largely recapitulated the ISG15 findings, with UCHL3^-/-^ cells showing reduced responsiveness to interferon stimulation. Interestingly, the magnitude of this effect was somewhat less pronounced for IRF7, suggesting that different ISGs may have varying dependencies on UCHL3 for optimal expression, reflecting the complexity of interferon-mediated transcriptional programmes.

We also analysed CXCL10, a chemokine that serves as both an interferon-stimulated gene and an important mediator of antiviral immunity (**Figure 5C**). Interestingly, the pattern of CXCL10 expression revealed the opposite effect, with UCHL3^-/-^ cells showing enhanced expression during viral infection but reduced responsiveness to interferon treatment. This paradoxical increase in CXCL10 during infection in UCHL3-deficient cells may reflect compensatory inflammatory responses or altered regulation of this particular gene in the absence of UCHL3.

To test whether this immunomodulatory function accounts for UCHL3’s pro-viral effects, we measured viral titres under conditions designed to equalise interferon responses between wild-type and UCHL3^-/-^ cells (**Figure 5D**). In the absence of exogenous interferon, UCHL3^-/-^ cells showed the expected reduction in Zika virus titres compared to wild-type controls, consistent with our previous findings. Crucially, when both cell types were treated with exogenous interferon, thus bypassing differences in endogenous interferon responsiveness, UCHL3^-/-^ cells continued to show significantly reduced viral titres compared to wild-type cells. This finding is further reinforced by direct comparison of interferon-treated wild-type and UCHL3^-/-^ cells (**Figure 5E**). Even under conditions where both cell types are exposed to identical interferon stimulation, UCHL3^-/-^ cells supported significantly lower levels of viral replication at both time points examined. The persistence of this phenotype under interferon-equalised conditions demonstrates that UCHL3’s pro-viral function operates through mechanisms that are fundamentally independent of its role in interferon signalling.

These data establish UCHL3 as a multifunctional protein with distinct roles in both immune signalling and viral replication. While UCHL3 functions as a positive regulator of interferon responses, its critical pro-viral effects during flavivirus infection operate through alternative mechanisms that are independent of interferon pathway modulation, potentially via viral RNA metabolism and condensate biology as the primary basis for its role in flavivirus pathogenesis.

## Discussion

Our findings establish UCHL3 as a key deubiquitylase that flaviviruses exploit to maintain their genomic stability through control of RNA-protein condensate dynamics. Cellular RNA metabolism occurs within highly organised, membrane-free biomolecular condensates^23,24^. These structures, including P-bodies^25^, stress granules^26^, and the newly characterised RNase L-induced bodies (RLBs)^9,27,28^, represent competing fates for viral RNA molecules, with dramatically different consequences for viral persistence. The central insight from our work is that UCHL3 functions as a molecular switch that biases sfRNA toward protective, pro-viral condensates rather than degradative, antiviral ones. This represents a previously unrecognised layer of post-translational control over RNA granule fate determination via ubiquitylation.

Based on our experimental evidence, UCHL3 appears to operate through distinct processes during flavivirus infection. The primary mechanism underlying UCHL3’s pro-viral activity centres on its role in maintaining sfRNA within protective P-body-like condensates. Here sfRNA can antagonise XRN1-mediated RNA decay and recruit host RNA-binding proteins that support viral replication. It is possible that UCHL3 deubiquitylates one or more critical components of sfRNA-containing condensates, such as G3BP1^29^, DCP1^30^, EDC3^30,31^, and UBAP2L^32^, thereby preventing their remodelling or dissolution. In UCHL3^-/-^ cells, these proteins likely become hyperubiquitylated, triggering condensate disassembly and allowing sfRNA to redistribute to RNase L-induced bodies where viral RNA decay is favoured. This model is well-aligned with the growing literature demonstrating that ubiquitin signalling shapes RNA granule dynamics. Other deubiquitylases, such as OTUD4, have been shown to regulate stress granule morphology and persistence through similar mechanisms^33^.

Emerging evidence suggests that sfRNA may serve an additional critical function in viral replication organelle (vRO) biogenesis^34,35^. Beyond simply directing RNA-protein condensates, viral 3’ UTR sequences can actively contribute to replication organelle formation. Multiple lines of evidence emphasise that vRO formation emerges from coordinated actions of viral non-structural proteins, host lipids, RNA-binding proteins, and the viral RNA itself^36–39^. It is possible that sfRNA nucleates or stabilises ribonucleoprotein platforms that couple to membrane-shaping factors to promote vRO biogenesis. This idea aligns with recent evidence that RNA-binding proteins can be hijacked to control vRO formation and viral RNA neosynthesis, such as IGF2BP2 in Zika virus infection^40^, and with broader Flaviviridae precedents where 3’-UTR-RBP interactions modulate lipid/ER programmes linked to replication membranes^41^.

While our data support the condensate-centric model outlined above, alternative possibilities exist. UCHL3 could influence the OAS–RNase L pathway more directly^42^, by deubiquitylating components of this pathway, rather than condensate scaffolds. Under this model, the observed changes in sfRNA localisation would represent secondary consequences of altered RNase L kinetics rather than the primary mechanism. In addition, our experiments were performed in transformed cell lines, which may not fully recapitulate the physiological responses of primary cells or intact tissues. Our future work will identify the specific proteins that UCHL3 deubiquitylates within sfRNA condensates, validated through in vitro deubiquitylation assays and rescue experiments with catalytically dead UCHL3 mutants to establish causality and distinguish enzymatic from scaffolding functions. Moreover, understanding UCHL3’s mechanism requires identifying the ubiquitin ligases that carry out the reverse reaction.

Our findings contribute to several evolving concepts in virology and cellular RNA biology. The demonstration that post-translational modifications can serve as molecular switches determining RNA fate represents a fundamental principle that likely extends beyond flavivirus infections. Recent studies have shown that a range of RNA viruses generate stable subgenomic species, including coronaviruses and togaviruses^43^. These pathogens may exploit similar mechanisms, suggesting that UCHL3 or related deubiquitylases could represent universal targets for RNA virus control.

## Acknowledgements

The Sanyal lab is supported by the Wellcome Trust (grants 220776/Z/20/Z and 223107/Z/21/Z to SS) and BBSRC (BB/Y000307/1 to SS).

## Methods

### Cell lines

A549, A549 UCHL3^-/-^, Huh7, and 293 cells were maintained in Dulbecco’s Modified Eagle Medium (DMEM) (Gibco) supplemented with 10% fetal bovine serum (FBS) (Gibco) and 20mM HEPES buffer (Sigma), at 37°C with 10% CO2. Vero cells were maintained in DMEM supplemented with 1% FBS and 20mM HEPES buffer, at 37°C with 10% CO2. Cells were passaged sub-confluently with 0.25% Trypsin-EDTA (Gibco).

### Virus stocks

Vero cells were infected with ZIKV strain MR-766 and incubated for 1h in FBS-free DMEM at 37°C with 10% CO2. Cells were washed with Dulbecco’s phosphate-buffered saline (PBS) (Gibco) and incubated in cDMEM for 5 days. Cell supernatants were collected, filtered, and aliquoted into sterile microtubes. Aliquots were stored at -80°C. Viral titres were determined using plaque assay.

### Virus infection

Cells were infected with ZIKV strain MR-766 or DENV2 (16681) at 0.5 multiplicity of infection (MOI), unless stated otherwise, and incubated for 1h in FBS-free DMEM at 37°C with 10% CO2. Cells were washed with PBS and incubated in cDMEM at 37°C with 10% CO2. Cell supernatants were harvested at the specified time points, and viral titres were determined using plaque assay.

### Plaque assay

Harvested supernatants were 10-fold serially diluted in FBS-free DMEM and incubated with 5×10^5^/ml Vero cells (≥80% confluency) in 24-well plates for 2h at 37 °C with 10% CO2. An overlay of 3% carboxymethyl cellulose (CMC) (Sigma) cDMEM was then added. After 84h of incubation, the CMC overlay was aspirated, and cells were washed with PBS. Cells were fixed and stained with Amido black (1g Naphthol blue-black powder, 60mL glacial acetic acid, 13.6g anhydrous sodium acetate in 1L demineralised water) (Sigma) for 1h, for quantification of plaque-forming units (PFU). Samples were measured in technical triplicates.

### Plasmid transfection

The Flag-HA-UCHL3 (#22564), BASU (#107250), and RNA motif plasmid cloning backbone (#107253) plasmids were obtained from Addgene. To produce the ZIKV sfRNA plasmid, sfRNA from ZIKV strain MR-766 was amplified using primers BSMBI sfRNA ZIKV Fw (5′-CAAGCTTGGAGACGGCACCAATCTTAGTGTTG-3′) and Rv (5′-AAGAGCTAGAGACGAGAAACCATGGATTTCC-3′) (**Table 1**). The resulting amplicon and a control RNA motif plasmid cloning backbone were digested with the BsmBI-v2 restriction enzyme (NEB) for 1h at 55°C. The digested fragments were purified and ligated using a DNA ligation kit (TaKaRa) at 4°C overnight. The recombinant ZIKV sfRNA plasmid was confirmed using DNA sequencing.

**Table 1:**
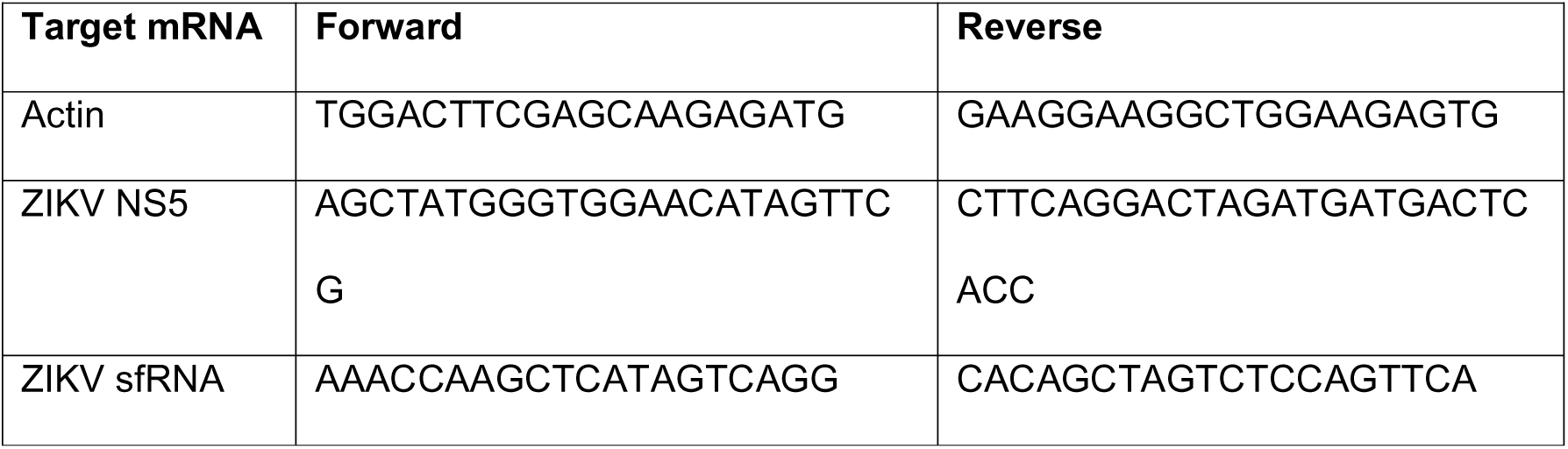
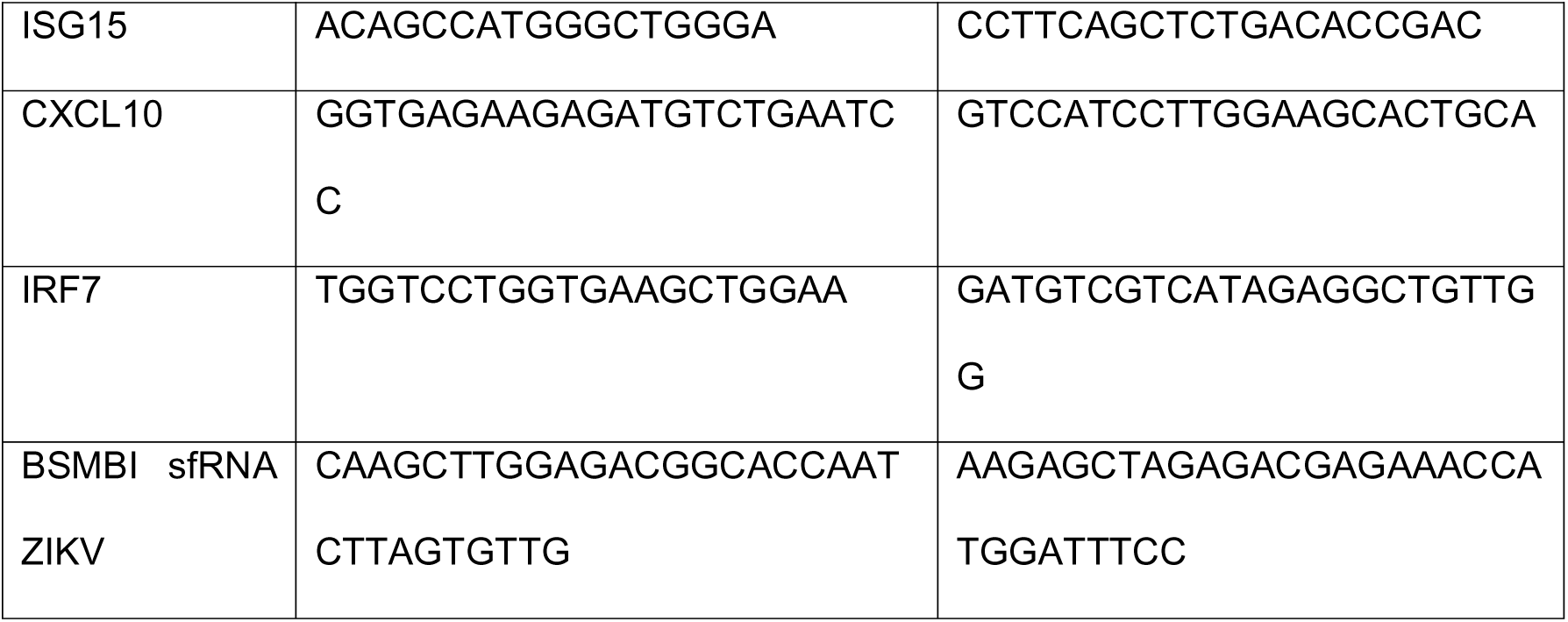
Primers

293 cells were seeded at 2×10^5^ cells/well (≥80% confluency) in 12-well plates in cDMEM and incubated for 24h at 37°C with 10% CO2. 2.5µg plasmid DNA was added to 250µL opti-MEM I Reduced-Serum Medium (Mirus). The diluted DNA mixture was added to *Trans*IT-LT1 reagent (Mirus) in a 1:2 ratio. The *Trans*IT-LT1 reagent:DNA complexes were incubated for 15 minutes at room temperature, then distributed dropwise to the seeded cells. 24h after transfection, cells were infected with mock (FBS-free DMEM) or ZIKV (MOI=0.5) for 24h or 48h; at these time points, cells were harvested, washed with PBS, and analysed.

### Interferon treatment

A549 and A549 UCHL3^-/-^ cells were seeded at 2×10^5^ cells/well (≥80% confluency) in 12-well plates and incubated in cDMEM for 24h at 37°C with 10% CO2. Cells were infected with mock (FBS-free DMEM) or ZIKV (MOI=0.5). 6h post-infection, cells were treated with 1000U/mL universal type-I IFN alpha (PBL Assay Science #11200-1) in cDMEM, for 24h and 48h; at these time points, cells were harvested, washed with PBS, and analysed.

### RT-qPCR

Harvested cells were lysed with RLT buffer (QIAGEN) for RNA analysis. Total RNA was extracted using the RNeasy® Mini Kit (QIAGEN). RNA concentrations were quantified using Nanodrop™ DS-11 FX (DeNovix). RNA samples were diluted in demineralised water to yield a final concentration of 50ng/µL. RNA samples were prepared using the One-Step TB Green® PrimeScript™ RT-qPCR Kit (TaKaRa Bio). The Master Mix was made following the manufacturer’s protocol, comprising 5µL RT-PCR Buffer III, 0.4µL TaKaRa Ex Taq® HS, 0.4µL PrimeScript RT Enzyme Mix II, 2.2µL RNase Free dH2O, and ROX Reference Dye 50X 0.2µL. The specified primers (Table 1) (Invitrogen) were diluted in the Master Mix. 9µL of Master Mix and 1.5µL of RNA sample were added per well in a MicroAmp® Fast Optical 96-Well Reaction Plate (Thermo Scientific). RNA transcripts were quantified using the ΔCT method, with actin as a reference, using the Applied Biosystems QuantStudio 3 Real-Time PCR System (Thermo Scientific). Samples were measured in technical triplicates.

### Immunofluorescence assay

A549 and A549 UCHL3^-/-^ cells were seeded at 5×10^4^ cells/well (≥80% confluency) on 24-well glass coverslips and incubated in cDMEM for 24h at 37°C with 10% CO2. Cells were infected with mock (FBS-free DMEM) or ZIKV (MOI=1) for 24h. cDMEM was aspirated, and cells were washed with PBS. Cells were fixed with 4% paraformaldehyde (PFA) in PBS for 15 minutes at room temperature. PFA was aspirated, and cells were washed 3 times with PBS. Cells were permeabilised with 0.1% Triton X-100 (Sigma) in PBS for 10 minutes at room temperature. 0.1% Triton X-100/PBS was aspirated, and cells were washed with PBS. Cells were blocked with 2% BSA/PBS for 30 minutes at room temperature. Cells were probed with the specified primary antibodies diluted at 1:100 in 2% BSA/PBS, incubated overnight at 4°C. Cells were washed with PBS 3 times, each for 5 minutes. Cells were incubated in the dark with Alexa Fluor™ 555 goat anti-mouse or 488 goat anti-rabbit secondary antibodies (Table 1) for 1h at room temperature, diluted at 1:500 in 2% BSA/PBS. Cells were washed with PBS 3 times, each for 5 minutes. Coverslips were mounted onto glass slides using 6µL of Duolink® in situ mounting medium with DAPI (Sigma) and exposed using a ZEISS LSM 880 confocal with Airyscan microscope (ZEISS). Images were analysed using ZEISS ZEN Microscopy Software (ZEISS), and fluorescent signal intensities were quantified using Image J.

### Immunoprecipitation assay

293 cells were seeded at 8×10^5^ cells/well (≥80% confluency) in 60mm plates and incubated in cDMEM for 24h at 37°C with 10% CO2. Cells were transfected with Flag-HA-UCHL3 plasmid or pcDNA3.1 (empty)-TAG control plasmid for 24h. Cells were infected with mock (FBS-free DMEM) or ZIKV (MOI=1) and incubated for 48h. Cells were lysed using Lysis Buffer (50mM HEPES pH 7.4, 150mM NaCl, 1mM MgCl2, 1% Triton X-100). Cell lysates were cleared by centrifugation at 13,000 rpm for 10 minutes at 4°C. Cleared lysates were incubated with Anti-FLAG® M2 Magnetic Beads (Sigma), with rotation for 3 hours at 4°C. The beads were washed 4 times in Lysis Buffer and resuspended in Sample Buffer. The resuspended beads were boiled for 10 minutes at 90°C. FLAG-tagged UCHL3 protein and its interactors were analysed by Western Blot.

### sfRNA-interactome capture assay

A modified version of RNA-protein interaction assay was employed to identify potential sfRNA-interacting proteins^44^. ZIKV sfRNA flanked by BoxB stem loops are specifically recognised by a λN peptide fused to the N-terminus of the BASU biotin ligase. The BoxB stem loops recruit the BASU fusion protein to the RNA, enabling biotinylation of proteins that interact with the sfRNA. The biotinylated proteins are isolated using streptavidin-conjugated beads^44^.

293 cells were seeded at 8×10^5^ cells/well (≥80% confluency) in 60mm plates and incubated in cDMEM for 24h at 37°C with 10% CO2. Cells were transfected with ZIKV sfRNA plasmid or control RNA motif plasmid cloning backbone, with BASU, for 24h. Cells were infected with mock (FBS-free DMEM) or ZIKV (MOI=1) and incubated for 24h. Cells were incubated with 200µM of biotin (Sigma) for 1h. Cells were lysed using Lysis Buffer. Cell lysates were cleared by centrifugation at 13,000 rpm for 10 minutes at 4°C. Cleared lysates were incubated with Pierce™ Streptavidin Magnetic Beads (Thermo Scientific), with rotation for 3 hours at 4°C. The beads were washed 4 times in Lysis Buffer and resuspended in Sample Buffer. The resuspended beads were boiled for 10 minutes at 90°C. Biotinylated proteins were analysed by Western Blot.

### RNase L activation assay

A549 wild-type (WT) and UCHL3^-/-^ cells were seeded 24 h before infection and inoculated with ZIKV (MR766 at MOI 2) for 1 h at 37 °C. At 24 h post-infection, cells were washed twice with ice-cold PBS and lysed on ice in 1× RNase-free lysis buffer (20–50 mM Tris-HCl, pH 7.5, 100–150 mM NaCl, 1–2 mM MgClLJ, 1 mM DTT) supplemented with protease inhibitor cocktail and RNase inhibitor. Lysates were clarified (14,000xg, 10 min, 4°C), protein-normalised (BCA), and maintained on ice. RNase L activity was quantified using a fluorogenic FRET RNA substrate in 96-well format as described before^45^. Clarified lysate was mixed with recombinant human RNase L and the dual-labelled RNA reporter in reaction buffer, and fluorescence was read kinetically at 37 °C 60 min in a plate reader. A standard curve was generated in parallel with serial dilutions of authentic (2’–5’)-pLJALJ. Assay conditions and data reduction followed established RNase L FRET protocols.

## Statistical analysis

Data are presented as the mean ± standard deviation (SD), unless stated otherwise. Statistical analyses were performed using GraphPad Prism 10. Significance was calculated using the unpaired Student’s two-tailed *t-*test, with Welch’s correction *t-*test where applicable, or two-way ANOVA with post-hoc Dunnett’s multiple comparisons test.

