## Supplemental information for "UCHL3 regulates subgenomic flaviviral RNA condensates to promote virus propagation"

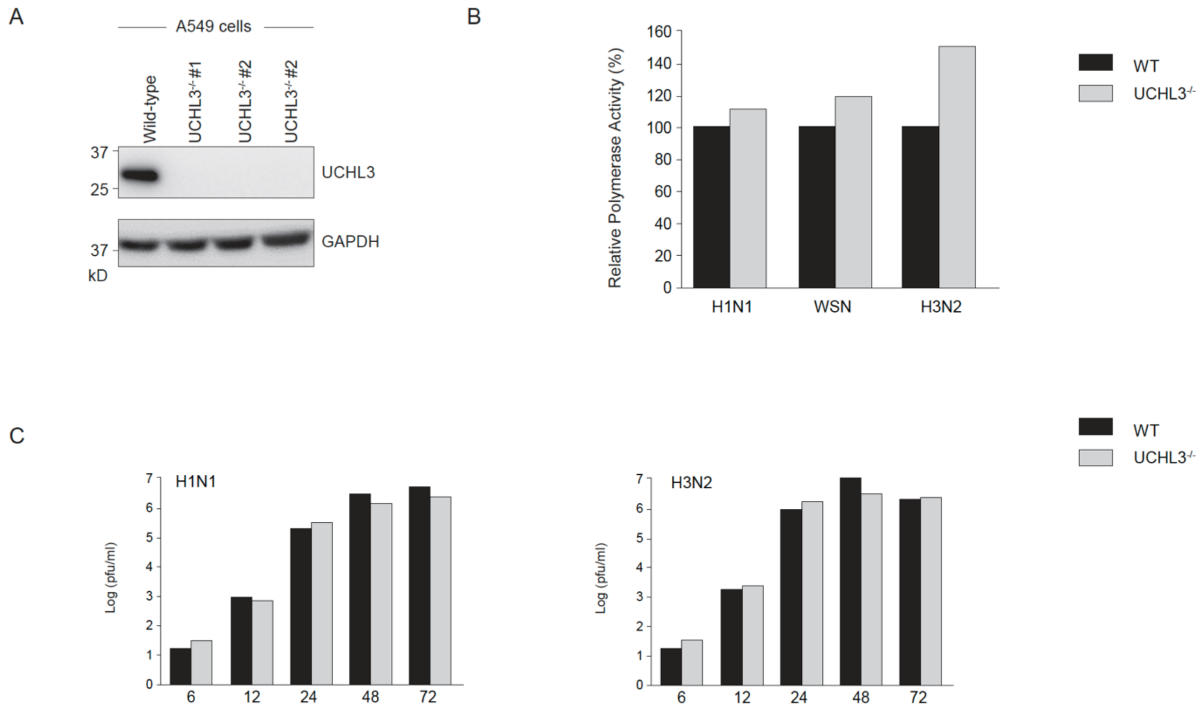


**Supplementary Figure S1. UCHL3 deficiency exhibits virus-specific effects on replication.**

**(A)** Immunoblot analysis of UCHL3 protein expression in three independent CRISPR-Cas9-generated A549 knockout clones ($UCHL3^{-/-}$ #1, #2, #2) compared to parental wild-type cells. GAPDH serves as loading control.

**(B)** Comparative analysis of viral polymerase activity demonstrates that $UCHL3^{-/-}$ cells exhibit increased polymerase function relative to wild-type (WT) controls across multiple influenza A virus strains (H1N1, WSN, H3N2). Relative polymerase activity is expressed as percentage of wild-type control levels.

**(C)** Time-course analysis of infectious viral particle production between WT and $UCHL3^{-/-}$ cells for both H1N1 (left panel) and H3N2 (right panel) influenza strains. Viral titres are expressed as pfu/ml across a 72-hour infection time course (6, 12, 24, 48, and 72 hours post-infection).

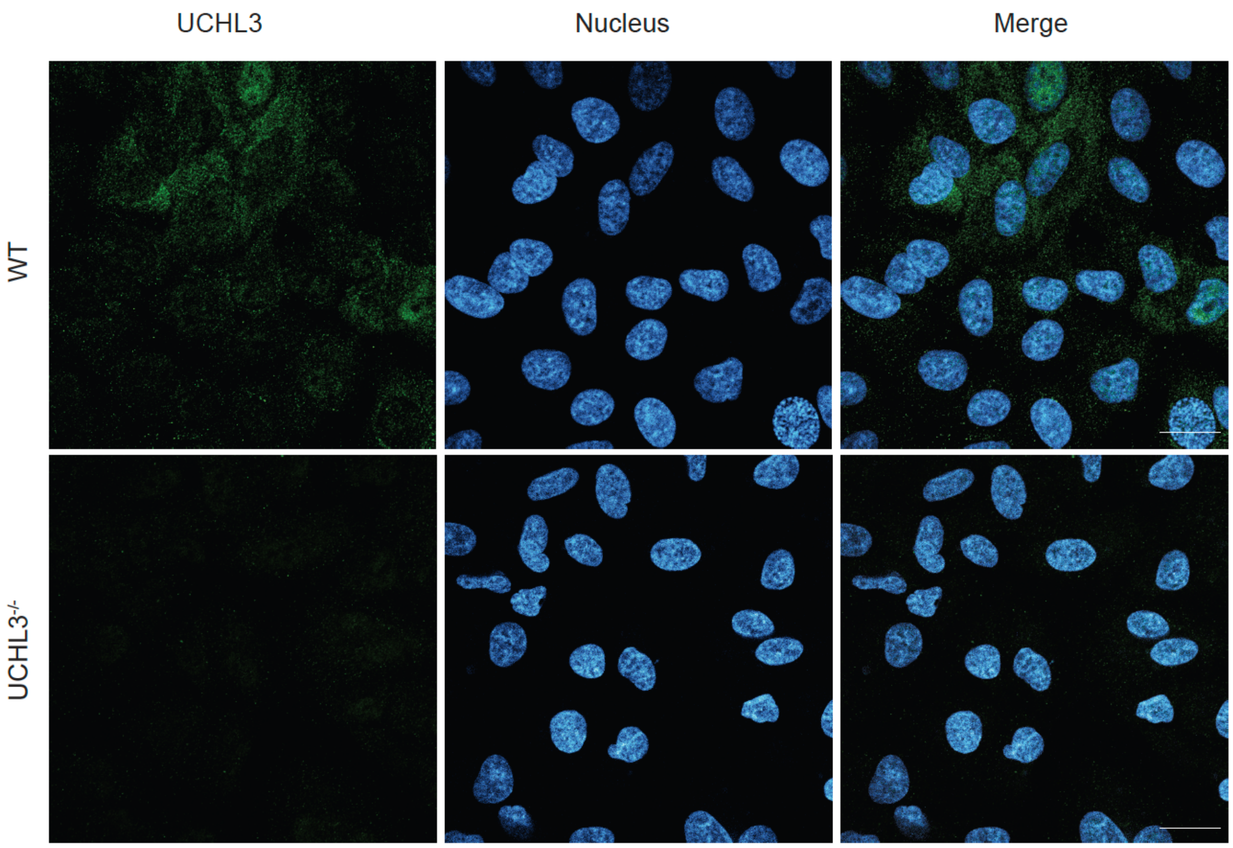


**Supplementary Figure S2. Validation of UCHL3 antibody specificity in knockout cell lines.**

Immunofluorescence analysis demonstrates antibody specificity and confirms complete UCHL3 ablation in CRISPR-Cas9-generated knockout cells. Representative confocal microscopy images of wild-type (WT, upper panel) and UCHL3$^{-/-}$ (lower panel) A549 cells stained for UCHL3 (green) with nuclei counterstained using DAPI (blue). Scale bar represents 20 μm.

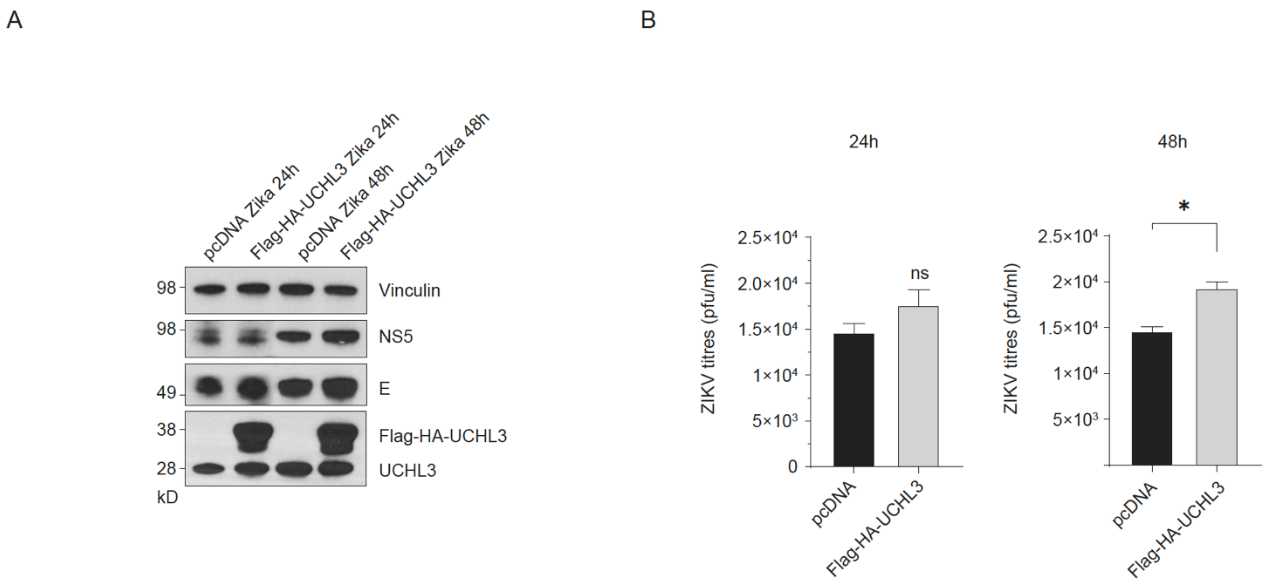


**Supplementary Figure S3. Ectopic UCHL3 expression enhances flavivirus replication and viral protein accumulation.**

**(A)** Immunoblot analysis of 293 cells transfected with pcDNA3.1 empty vector control or Flag-HA-UCHL3 expression construct and subsequently infected with ZIKV at 0.5 MOI. Protein expression was assessed at 24 and 48 hours post-infection. Vinculin serves as loading control. Viral non-structural protein NS5 and structural envelope protein E accumulation in Flag-HA-UCHL3-expressing cells compared to vector controls, measured at 24 and 48 hours post-infection. Endogenous UCHL3 levels remain detectable across all conditions, providing internal validation of antibody specificity and baseline protein expression.

**(B)** Viral titres were determined using plaque assay in Flag-HA-UCHL3-transfected cells relative to pcDNA3.1 control transfections at 24 and 48 hours post-infection. Data represent mean ± SD from 3 independent experiments. *$P < 0.05$.
